# Identification of human RNA Polymerase II interactors at early stages of transcription

**DOI:** 10.1101/2025.10.08.681243

**Authors:** Abdullah Ozer, Jin Joo Kang, Yuliang Tang, Kumar Yugandhar, Sun Yu, Michael D. DeBerardine, Karim Omar, John T. Lis, Haiyuan Yu

## Abstract

RNA Polymerase II (Pol II) transcription is highly regulated at two early steps in the transcription cycle: <u>P</u>re-Initiation <u>C</u>omplex (PIC) assembly with its coupled initiation, and promoter-proximal pausing with its controlled release. Here, we developed an optimized biochemical purification method that captures endogenously tagged chromatin-bound Pol II complexes under native conditions at these rate-limiting steps. We then identified a large set of Pol II interactors by mass spectrometry and determined the footprints of these assemblies on promoters with high resolution. Many well-known and new or understudied factors were identified as associated with the PIC and promoter-proximal paused complexes, indicating that despite decades of efforts, these rate-limiting steps of the transcription cycle are far from being completely understood. The new and understudied factors implicate novel mechanisms of regulation that will need to be characterized to fully understand Pol II regulation.

## Introduction

RNA Polymerase II (Pol II) transcription is a highly regulated process that is responsible for the production of all mRNAs and many other RNAs in eukaryotic cells. This transcription is tightly regulated by chromatin-associated protein networks that act on distinct stages of the transcription cycle^1,2^. Pol II undergoes numerous transformations and dramatic changes in molecular interactions as it progresses through the steps of the transcription cycle. Among these steps, the pre-initiation complex (PIC) formation and transcriptional pausing represent two major regulatory checkpoints. These steps are mediated by dynamic and coordinated interactions involving transcription factors, cofactors, and chromatin-modifying complexes^3–6^. Structural studies have highlighted the roles of General Transcription Factors (GTFs) and Mediator, and also DRB-sensitivity Inducing Factor (DSIF) and Negative Elongation Factor (NELF) complexes^7^, in orchestrating the assembly and stabilization of Pol II during PIC formation and promoter-proximal pausing, respectively^5,8–10^. The release of paused Pol II into productive elongation for essentially all genes is regulated by the activity of P-TEFb kinase^11,12^, whose recruitment to genes is regulated by TFs and cofactors (reviewed in Refs^13,14^).

Many of the factors involved in transcription and its regulation including the Pol II machinery are large molecular complexes (LMCs). Pol II is composed of 12 subunits, where the largest two subunits (POLR2A/Rpb1and POLR2B/Rpb2) form the active site. Rpb1’s C-terminal domain (CTD) domain is composed of 52 repeats of a heptapeptide consensus (YSPTSPS) functions as a post-translation-modifiable scaffold for LMC interactions^15^ and a target for multiple kinases and phosphatases^16^, which themselves are often part of LMCs that regulate Pol II’s transitions through steps in the transcription cycle. Pol II’s recruitment to promoters and its initiation is guided by General Transcription factors (GTFs) including TFIIA, IIB, IID, IIE, IIF, and IIH that combine with Pol II to create a PIC. The efficiency of this recruitment is usually often controlled by collections of transcription factors (TFs) that bind specific DNA sequence motifs of promoters as well as specific subunits of the 30+ subunit, Mediator complex that also interacts tightly with RNA Pol II^17^. These protein interactions thereby create large multi-docking Mediator hubs where TFs recruit Pol II and provide the specificity of gene regulation. Once the PIC is formed, it rapidly initiates transcription^18^ and moves to the promoter-proximal pause site where pausing is stabilized by molecular complexes of DSIF, which is composed of Spt5 and Spt4, and its interacting complex NELF, which is composed of NELF-A, -B, C/D, and -E. The release of Pol II to productive elongation also requires participation of specific TFs and cofactors most notably, P-TEFb kinase or its larger Super Elongation Complex (SEC) that phosphorylate multiple components of the promoter-proximal paused Pol II complex. The LMCs that control these steps and subsequent steps of the Pol II transcription cycle remain challenging to characterize because of their large size, subunit complexity, varying stability of their interactions, and numerous interactions that execute specialized functions on diversely-regulated sets of genes.

In addition to the earlier biochemical and genetic studies, more recent structural work revealed substantial information about Pol II and its interactors: basic form of Pol II from the magnificent X-ray crystallography by the Kornberg and Cramer Labs^19,20^. More recently, spectacular cryo-EM derived structures by the Cramer lab of Pol II assembled with other LMCs and particular factors have provided models for Pol II promoter-proximal pausing and its transition to an elongation state^5,6,5,21,22^. However, *in vitro* assembly of Pol II complexes from purified complexes can only include known factors, and inevitably involves making assumptions about assembly conditions and artificial DNA/RNA hybrid templates that may be biased by our incomplete understanding of Pol II and associated LMCs.

The study of transcription regulation has also benefited from mammoth-scale efforts like ENCODE that provided information about the binding of specific TFs and the co-occupancy with individual LMC subunits (e.g., P300, MED1) along the genome^23^, though they do not address the nature and function of LMCs that associate with those TFs. Likewise, the distribution of transcriptionally-engaged RNA Pol II have been tracked sensitively and at base-pair resolution across genomes by nascent transcriptomics methods like PRO-seq (reviewed in Ref^24^). Despite these advances, our understanding of the full composition and regulation of chromatin-associated Pol II interactomes at the PIC and paused states remains strikingly incomplete. This gap stems from limitations in existing methodologies that fail to preserve interactions specific to transcriptionally engaged chromatin at particular steps in the transcription cycle^25–27^.

Although ChIP-seq and related techniques have been quite successful in identifying chromosomal regions bound by bait factors, it is not appropriate for finding other protein interactors associated with the bait, because bait proteins are purified from whole cell lysates after cross-linking and sonication in canonical ChIP protocols^28^. This is especially problematic for Pol II because Pol II is highly abundant (∼320,000 Pol II molecules/human cell) and ∼50% of them are not bound on chromatin (thus not active)^29^.

In this study, we employed a high-resolution approach to map the native, chromatin-associated interactome of Pol II during the PIC and Pause stages. We generated a CRISPR-Cas9-engineered human cell line expressing N-terminal GFP-tagged RPB1 at endogenous levels, enabling the isolation of Pol II complexes under physiological conditions. Using this system, we successfully purified N-terminal GFP-tagged RPB1 in both the pre-initiation complex—as supported by structural investigations highlighting the coordination of TFIIH and Mediator in transcription initiation^3^—and in the paused state of the transcription cycle, utilizing EnChAMP-MS (<u>En</u>dogenous <u>Ch</u>romatin-<u>A</u>ssociated <u>M</u>acromolecular complex <u>P</u>urification followed by <u>M</u>ass <u>S</u>pectrometry) and EnChAMP-Seq (followed by DNA <u>seq</u>uencing). These techniques allowed us to isolate and characterize Pol II-associated complexes under native conditions, preserving chromatin-bound interactions while minimizing artifacts. Compared with other published studies using whole cell lysates^30,31^, our EnChAMP-MS of GFP-RPB1 identified significantly more known Pol II interacting proteins and complexes.

## Results

### EnChAMP: an Enhanced Chromatin-Associated Macromolecule Purification workflow

To investigate the transcriptional behavior of RNA Polymerase II (Pol II), we engineered an HCT116 cell line expressing N-terminally GFP-tagged POLR2A/RPB1 (hereon referred to as GFP-RPB1) from the endogenous RPB1 locus (**Fig. 1A**). Homozygous genomic integration and proper processing of engineered GFP-RPB1 gene was confirmed by genotyping and Western Blotting. A control cell line expressing GFP alone (GFP control) was also established by including a P2A ribosomal skip sequence between the GFP and the RPB1 at the native RPB1 locus. These designs ensure normal levels of GFP-RPB1 expression compared to RPB1 in the parental HCT116 cell line, as well as comparable expression levels of GFP in the control line, allowing a direct comparison of complexes purified from GFP-RPB1 and GFP control cell lines. Normal growth rate observed for these cell lines ensures the integrated GFP does not significantly alter the function of the tagged protein and transcription in general.

**Figure 1.**
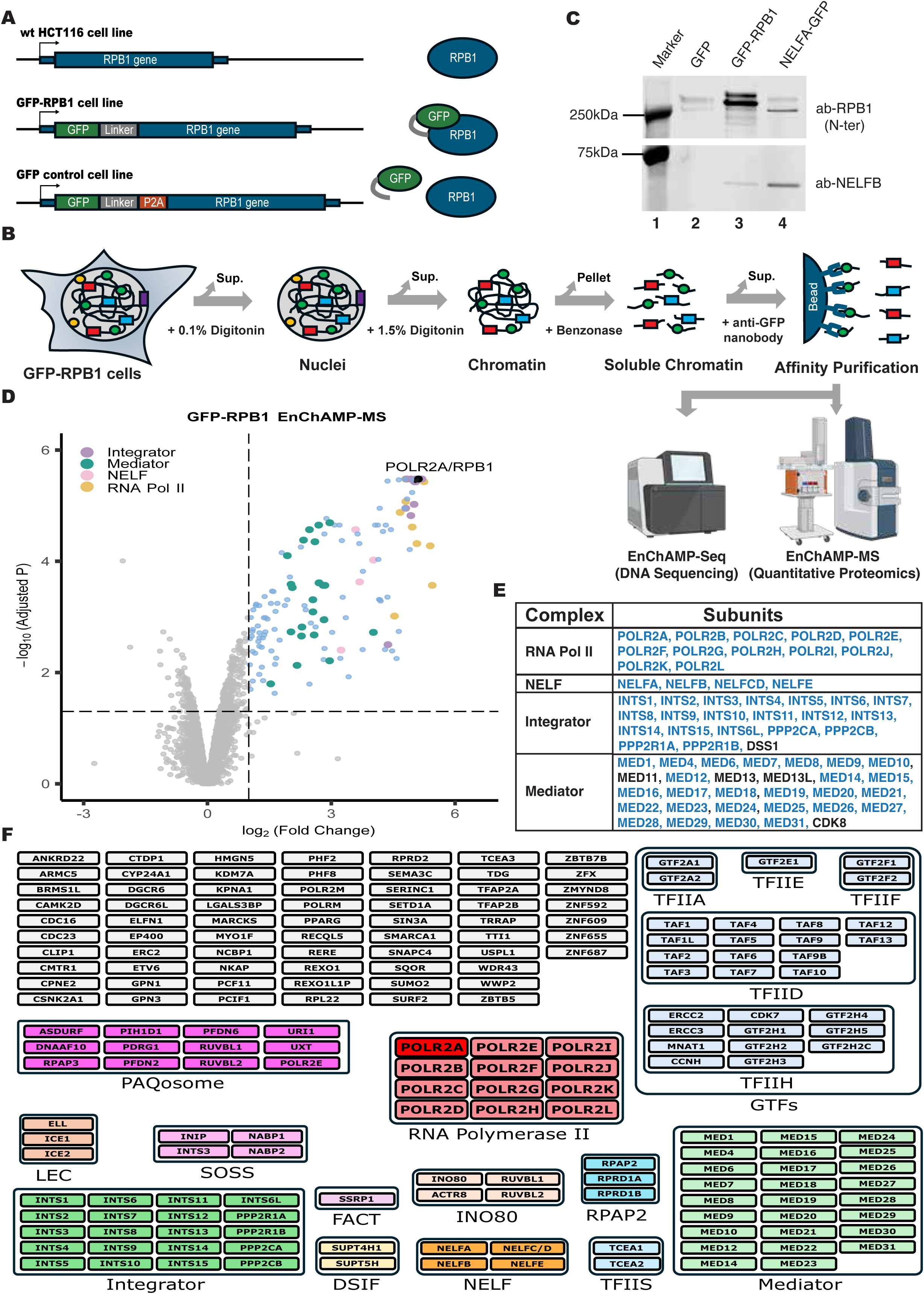
Enhanced Chromatin Associated Macromolecule Purification (EnChAMP) of Human RNA Polymerase II. A) HCT116 parental cell line was engineered with CRISPR/Cas9 system to introduce an in-frame GFP or GFP-P2A sequence at the start codon of the RPB1 gene to produce GFP-RPB1 fusion or uncoupled GFP-P2A and RPB1 proteins in GFP-RPB1 and GFP control cell lines, respectively. For the NELFA-GFP cell line expressing a C-terminally GFP tagged NELF-A fusion protein, the GFP coding sequence with a short linker was inserted in-frame with the NELF-A gene immediately upstream of the stop codon. All engineered cell lines were verified to be homozygous, and viable with normal growth rates. B) Schematic depiction of the major steps of EnChAMP including nuclei isolation, chromatin isolation, chromatin solubilization, and anti-GFP nanobody affinity purification. Fractions discarded were indicated with curved arrows. Resulting EnChAMP material used for Quantitative Proteomics (EnChAMP-MS) for protein identification and for DNA Sequencing (EnChAMP-Seq) for determining the genomic location of the purified complexes. C) A representative Western Blot analysis of EnChAMP samples from GFP control, GFP-RPB1, and NELFA-GFP cells using RPB1 and NELF-B specific antibodies. D) A representative Volcano plot of protein enrichments in GFP-RPB1 over GFP control cells by EnChAMP-MS. Log2(Fold change) and −log10(Adjusted p-value) values are plotted in x- and y-axes. Positive and negative log2(Fold Change) values indicate enrichment in GFP-RPB1 and GFP control samples, respectively. E) RNA Pol II and several well-known RNA Pol II associated complexes and their subunit compositions are shown, where EnChAMP-MS detected GFP-RPB1 interactors are indicated with a blue font. F) Complete list of EnChAMP-MS identified GFP-RPB1 interactors are grouped based on the transcription related protein complexes that they belong to. A large number of factors that are not known to be subunits of well-known transcription related complexes are shown as individual grey colored boxes. POLR2A/RPB1 bait protein indicated with a red colored box.

To identify Pol II interactors that are functionally coupled to transcription, we optimized a multi-step biochemical purification procedure, which we named <u>En</u>dogenous <u>Ch</u>romatin-<u>A</u>ssociated <u>M</u>acromolecular complex <u>P</u>urification (EnChAMP), producing a soluble chromatin fraction that is then subjected to affinity purification by anti-GFP nanobody (**Fig. 1B**). EnChAMP begins with nuclei isolation in the presence of low concentration (0.5%) Digitonin that provides selective permeabilization of the plasma membrane while maintaining nuclear envelope integrity (**Supp.** Fig. 1A). This approach effectively minimized cytoplasmic contamination, resulting in intact nuclei with preserved protein-DNA complexes. The use of a high concentration (1.5%) of Digitonin in chromatin isolation buffers provided a precise and effective method for obtaining high-quality chromatin. Digitonin’s specificity, a non-ionic detergent targeting cholesterol containing membranes^32,33^, allowed selective permeabilization of nuclear membranes while preserving chromatin structure. The resulting chromatin pellet is digested with benzonase to obtain a soluble chromatin fraction free of insoluble chromatin debris, which is then subjected to affinity purification using an anti-GFP nanobody. The chromatin immunoprecipitation workflow included an optimized washing and elution protocol to enhance specificity and protein recovery. Sequential washes with chromatin isolation buffers effectively removed non-specific proteins and cellular debris, ensuring high-purity chromatin (**Supp.** Fig. 1A). Finally, the purified material is analyzed by mass spectrometry (EnChAMP-MS) to identify the composition of the Pol II associated complexes; the DNA component of these same isolated chromatin complexes is also analyzed by sequencing (EnChAMP-seq) that maps the genomic location of the isolated LMCs like Pol II, and thereby determines the major step in the transcriptional cycle in which these LMCs are captured.

To complement our findings with GFP-RPB1, we generated an additional endogenously GFP-tagged cell line for NELF-A, a subunit of the NELF complex (**Fig. 1A**). NELF, along with the DSIF complex, has well-established functions in Pol II pausing thus are known as pausing factors. Upon pause release, NELF dissociates from Pol II whereas DSIF gets transformed into an elongation factor maintaining its interactions with elongating Pol II. While striving for stringency, we optimized our EnChAMP protocol with respect to incubation conditions like buffer composition, time, and temperature to retain the weak interaction of Pol II with the NELF complex. Indeed, our optimized EnChAMP protocol is gentle enough to capture the weak Pol II-NELF complex interaction (**Fig. 1C**).

Our method ensures the integrity of chromatin interactions by employing a gentler enzymatic digestion to shear DNA and by digitonin-based permeabilization to selectively disrupt cellular membranes while preserving nuclear architecture^30^. This strategy involving mild buffer conditions and rapid processing enabled us to retain weak and transient protein interactions that are often lost with harsher methods like sonication or cross-linking. Our approach of using endogenously-tagged RPB1/NELF-A allowed the identification of their native protein interaction partners without artificial stabilization or complications due to overexpression, enhancing the specificity of chromatin-bound complex profiling. In contrast to large-scale interactome studies based on whole-cell lysates or proximity labeling^31,34^, our method captures the transcriptionally relevant subset of protein networks directly engaged with chromatin during defined transcriptional states. This methodological framework revealed novel protein-protein interactions and provided an improved, high-confidence map of regulatory protein networks and potential regulatory mechanisms associated with the PIC and paused Pol II states. These networks are essential for maintaining transcription fidelity, coordinating responses to cellular signals, and influencing chromatin architecture. Our findings provide high-confidence maps of protein interactions under native conditions. By enabling a more complete annotation of factors associated with transcriptional regulation and their precise genomic location, our study not only enhances the current understanding of transcription initiation and pausing but also lays the foundation for future studies of factors likely involved in the mechanism of transcriptional regulation and dysregulation in development and disease.

### EnChAMP-MS identifies a large set of factors associated with Pol II

Our initial pilot EnChAMP-MS study was done using TMT labeling with canonical Data-Dependent-Acquisition (DDA-TMT) quantitative proteomics workflow^35^. However, with the rapid development of Data-Independent-Acquisition (DIA) methods (enabled by the powerful deep learning based search algorithms^36,37^), new label free quantification (LFQ) methods have been shown to be even more powerful, providing much increased sensitivity and coverage^38,39^. We thus optimized a DIA-LFQ workflow to process our EnChAMP samples using Bruker timsTOF HT that led to identification and quantification of twice as many protein targets compared to DDA-TMT.

Using our DIA-LFQ-based EnChAMP-MS, we identified 185 proteins associated with Pol II by comparing EnChAMP samples from GFP-RPB1 cells to that of GFP control cells with a Fold Change cut-off of 2 and Adjusted p-value (FDR) cut-off of 0.05 (**Fig. 1D**). The bait protein, POLR2A/RPB1, is among the most significantly enriched proteins. Subunits of many well-characterized Pol II and associated regulatory complexes such as Integrator, NELF, and Mediator are also among the enriched proteins. Our approach successfully captured entire sets of RNA Pol II core, NELF, and Integrator subunits, and a nearly complete set of subunits for Mediator and several other regulatory complexes (**Fig. 1E** and **Supp. Table I**). We repeated the GFP-RPB1 EnChAMP-MS experiment 3 times each with 4 biological replicate samples of GFP-RPB1 and GFP control, and observed a substantial overlap between the identified Pol II interactors in each experiment with a reproducibly high enrichment across the three independent EnChAMP-MS experiments (**Supp.** Fig. 1B and 1C). Using these data, we generated a high confidence Pol II interactor list containing 185 proteins that showed significant interactions in 2 out of the 3 independent experiments (**Fig. 1F**). Notably, the Pol II interactors excluded from our high confidence list, that are unique to each experiment (total of 298 proteins), include many well-known transcription related proteins such as SUPT6H/SPT6, FACT complex subunit SUPT16H/SPT16, PAF complex subunit LEO1, Mediator subunit CDK8, BRD4, Super Elongation Complex subunit AFF4, and others (**Supp.** Fig. 1B and **Supp. Table I**). This further supports the stringency of our method and selection criteria implemented.

Further categorization of the high-confidence GFP-RPB1 enriched proteins revealed multiple protein complexes: some with long-standing functions in transcription, some that were recently implicated, but others with as yet little or no implicated functions in transcription (**Fig. 1F**). A large set of factors belonging to General Transcription Factors (GTFs) including TFIIA, TFIID, TFIIE, TFIIF, and TFIIH, subunits of TFIIS, RPAP2, INO80, DSIF, SOSS, LEC, and PAQosome complexes. GTFs are critical for recognition of promoter elements and proper recruitment and placement of Pol II for transcription initiation^40^. Mediator complex, together with GTFs, functions in assembly of PIC and acts as an integrator of transcriptional signals from sequence-specific transcription factors and enhancers to Pol II at gene promoters^17^. DSIF and NELF complexes are key pausing factors responsible for stabilization Pol II pause to ensure proper modification of Pol II for productive elongation following P-TEFb mediated phosphorylation of Pol II, DSIF, and NELF for pause release^10^. TFIIS (TCEA1 or TCEA2) rescues backtracked Pol II by triggering the cleavage of RNA at the active site thus realigning the Pol II active site with the 3’ end of the RNA^41^. Integrator complex can terminate Pol II that encounters impediments during early elongation and pausing^41^. While implicated in the cytoplasmic assembly of Pol II subunits and Pol II nuclear import, recent cryo-EM structures indicate that RPAP2 is expected to sterically inhibit PIC formation, creating a checkpoint for initiation^42,43^, and it also acts subsequently to initiation in the RPRD complex to inhibit Pol II CTD phosphorylation^44^. SOSS complex, harboring the INTS3 subunit of Integrator complex, promotes DNA repair as a single-strand DNA sensor at sites of double-strand breaks^45^. Little Elongation Complex (LEC) promotes transcription of small nuclear RNAs (snRNAs) by Pol II^46^. INO80 complex is an ATP-dependent chromatin remodeler with wide ranging functions in gene expression, DNA repair, and DNA replication^47^. The PAQosome is a 12-subunit chaperone complex^48^ involved in assembly of various protein complexes including Pol II^49^.

Among the other interactors identified in our dataset that are not classified as subunits of well-established protein complexes (gray colored individual boxes in **Figure 1F**), several of them have also been implicated, some more than others, in transcription regulation, for example, ARMC5, CMTR1, CTDP1, GPN1/GPN3, PCF11, and RECQL5. ARMC5 interacts with the ubiquitin ligase CRL3 to terminate excessive or defective RNA Pol II molecules at the early stages of the transcription cycle^50,51^. CMTR1 is an mRNA capping, adds methyl to 2’OH of the first nucleotide in the Cap^52^. CTDP1, also called FCP1 phosphatase, is critical for dephosphorylating and recycling Pol II^53^ and for dissociation of capping enzymes from the elongation complex^54^. PCF11 is a component of pre-mRNA cleavage complex II, which promotes transcription termination by RNA Pol II ^55^. RECQL5 is a DNA Helicase known to bind Pol II^56,57^.

To evaluate the performance of our EnChAMP-MS result of GFP-RPB1, we compared our results to 3 previously published proteomic studies of Pol II: two of which, BioPlex^31^ and OpenCell^30^, purified RPB1 from whole cell lysates (**Supp.** Fig. 2A-D). OpenCell and BioPlex yielded purification of very few known transcription-related complexes and these are incomplete, including RNA Pol II complexes missing several subunits. In contrast, our EnChAMP-MS identified all 12 core subunits of Pol II, together with many other known factors and complexes. These results highlight the importance of chromatin isolation, rather than simply using whole cell lysate, to identify proteins involved in transcription using proteomics, because it has been estimated that about half of Pol II complexes in the cell are not bound on the chromatin, and thus not active engaged in transcription^29^. The third published proteomic study of Pol II used an overexpressed HA-RPB3 for chromatin purification. As expected, their data are indeed more comprehensive than OpenCell^30^ and BioPlex^31^. However, they still detected many fewer known interactors than our EnChAMP-MS. We carefully compiled a list of 38 known complexes with 247 proteins in total. The HA-RPB3 study detected 16 complexes; while EnChAMP-MS detected all 16, plus 11 more complexes. For these complexes, EnChAMP-MS on average detected 40% of the subunits of each complex, much more complete than the HA-RPB3 study (only 6%; **Supp.** Fig. 2E). The comparison with these published studies confirm the effectiveness of our EnChAMP-MS protocol in its use of endogenously tagged proteins, isolation efficiency, preservation of weaker interactions, MS workflow (DIA-LFQ), and instrument sensitivity (Bruker timsTOF HT) for comprehensive identification of proteins and LMCs associated with transcription.

### EnChAMP captures the RNA Pol II predominantly in the PIC and paused states

To selectively enrich Pol II in specific stages of transcription, we utilized drugs to inhibit transcriptional process at specific steps (**Fig. 2A**). The Pol II transcription cycle consists of PIC assembly leading to transcription initiation, pausing, elongation, termination, and finally recycling of Pol II and associated factors for additional rounds of transcription. Recycling takes place off of the chromatin, thus will not be captured by EnChAMP. NVP-2, a specific P-TEFb (CDK9/CYCT1 heterodimeric complex) inhibitor^58^, can effectively halt the release of paused Pol II into productive elongation, while already elongating Pol IIs continue transcription more or less unaffected. Given enough time (i.e., 1 – 2 hours) NVP-2 treatment^59^, much like CDK9 inhibitor Flavopridol^12,60^, would lead to near complete elimination of elongating Pol IIs from chromatin leaving only Pol II in pause and PIC. The Triptolide, a TFIIH helicase inhibitor, blocks transcription initiation, eliminating all paused and elongating Pol IIs from chromatin and just leaving Pol IIs at the PIC^12^.

**Figure 2.**
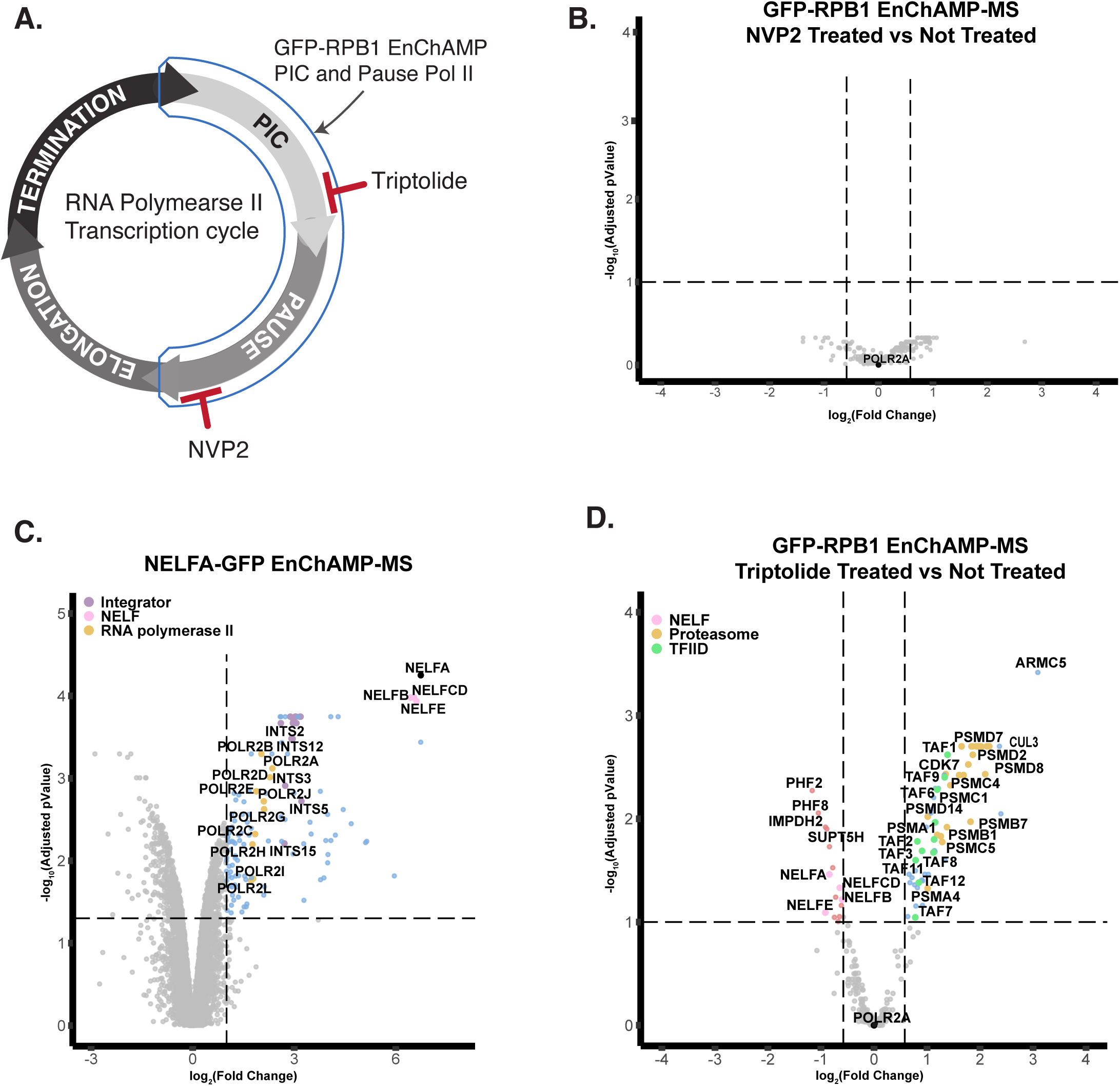
EnChAMP-MS analysis of RNA Pol II interactions at major regulatory steps of RNA Pol II transcription cycle – PIC assembly/initiation and Pol II pausing. A) Schematic of the RNA Pol II transcription cycle, which proceeds through PIC assembly and transcription initiation, Pol II pausing, productive elongation, termination, and recycling. PIC assembly/initiation and pausing represent the two major regulatory steps and initiation and pause release can be inhibited with Triptolide and NVP2, respectively. B) A representative volcano plot of GFP-RPB1 EnChAMP-MS comparing NVP2 treatment to untreated cells. Positive log2(Fold Change) values indicate enrichment in +NVP2 condition over -NVP2. This experiment was repeated 3 times, each with 4 replicates, with similar results. C) Volcano plot of NELFA-GFP EnChAMP-MS. Positive log2(Fold Change) values indicate enrichment in NELFA-GFP samples over GFP control samples. 4 biological replicates were analyzed in this experiment. D) A representative volcano plot of GFP-RPB1 EnChAMP-MS comparing Triptolide treatment to untreated cells. Positive log2(Fold Change) values indicate enrichment in +Triptolide condition over -Triptolide. This experiment was repeated 3 times, each with 4 replicates, with similar results.

Upon NVP-2 treatment we did not observe any significant difference in either enrichment or depletion in interactions of GFP-RPB1 using a Fold Change cut-off of 1.5 and Adjusted p-value (FDR) cut-off of 0.1 (**Fig. 2B**). Considering the fact that EnChAMP-MS of GFP-RPB1 readily detects many pause and PIC components and lacks subunits of elongation complexes such as PAF1 (**Fig. 1F**), we conclude EnChAMP predominantly captures factors associated with Pol II in early stages of transcription, the PIC and pause, whereas some factors associated with elongating Pol IIs appear underrepresented, like PAF1, perhaps due to its less stable association with Pol II^61^. Using a complementary NELFA-GFP cell line for EnChAMP-MS, we identified over eighty proteins associated with RNA Pol II pausing (**Fig. 2C** and **Supp. Table II**). Validating our method, we recovered the entire NELF complex, composed of NELFA, NELFB, NELFC/D, and NELFE, along with Integrator, and DSIF (**Supp. Table II**) with well-established roles in Pol II pausing at a subset of genes^62^. This overlap underscores their integrated functions within the transcriptional apparatus. However, GFP-RPB1 displayed a much broader interaction network, some of which is uniquely associated with PIC, such as the Mediator complex, and general transcription factors (e.g., TFIIA, TFIID, TFIIF, and TFIIH), and others.

### Triptolide treatment helps further dissect LMCs associated with PIC vs. paused Pol II complexes

Triptolide inhibits the XPB subunit of TFIIH, blocking its ATP-dependent DNA translocase activity^63^. Therefore, a short-term triptolide treatment (20 min at 1μM) allows us to specifically purify PIC^12^. As expected, interaction with GTFs increased, while interaction with pause factors such as the NELF complex and SPT5 subunit of DSIF decreased, using a Fold Change cut-off of 1.5 and Adjusted p-value (FDR) cut-off of 0.1 (**Fig. 2D**).

In contrast, transcription initiation and proteasomal regulation factors were up-regulated after triptolide treatment. The TFIID complex which plays a role in PIC formation showed increased enrichment. Also enriched are proteins whose association likely deal with defective Pol II initiation. These include ADRM1, ARMC5, CUL3, and BRD2. ARMC5 and CUL3 participate in transcriptional regulation and cellular stress responses^50,51,64^. ADRM1, a proteasomal ubiquitin receptor, regulates protein turnover and homeostasis^65^. Triptolide also increased the interaction with several proteasome subunits, including PSMA, PSMB, PSMC, and PSMD. BRD2, belonging to the bromodomain and extra-terminal (BET) family, facilitates chromatin remodeling and transcriptional activation by recognizing acetylated histones^66^. Its elevated enrichment may reflect either persistent association of triptolide-blocked Pol II with initiation-promoting factors or a compensatory mechanism that boosts initiation in response to inhibition.

### EnChAMP-seq results of GFP-RPB1 confirm enrichment of RNA Pol II in PIC and pause states

To gain insights about the genomic position and thus the transcription cycle stage of RNA Pol IIs captured by GFP-RPB1 EnChAMP, we extracted genomic DNA fragments that co-purified with Pol II and sequenced them with high-throughput nextGen sequencing (EnChAMP-seq) (**Fig. 1B**). First we looked at genomic fragments obtained in soluble chromatin fraction (Input) used for anti-GFP nanobody affinity purification in EnChAMP and found them to more or less uniformly cover the genome (**Supp.** Fig. 1D). GFP control EnChAMP samples show a weak but uniform background signal arising most likely from nucleosomal protection of genomic DNA (**Supp.** Fig. 1E). EnChAMP-seq fragments that we obtained from GFP-RPB1 samples revealed a robust enrichment of signal downstream of TSSs, consistent with promoter-proximal paused Pol II positioning (**Fig. 3A** and **Supp.** Fig. 1E). Visual inspection of mapped reads in genome browser revealed significant enrichment of EnChAMP-seq signal near gene TSSs (**Fig. 3A**). The vast majority of genes, like GAPDH, showed EnChAMP-seq signal at the TSS with minimal to background levels of signal in the genebody, beyond 500bp of TSS. Only a small fraction of genes, like EEF1A1, showed any appreciable genebody signal while the TSS signal was readily detectable. Both GAPDH and EEF1A1 are highly expressed as the genebody Pol II are readily detectable with ChIP-seq and PROseq (**Fig. 3A**). Importantly, the underrepresentation of elongating Pol II in EnChAMP-seq cannot simply be due to failure of transcriptional elongation, as RPB1 is homozygously tagged with GFP and the cells grow normally.

**Figure 3.**
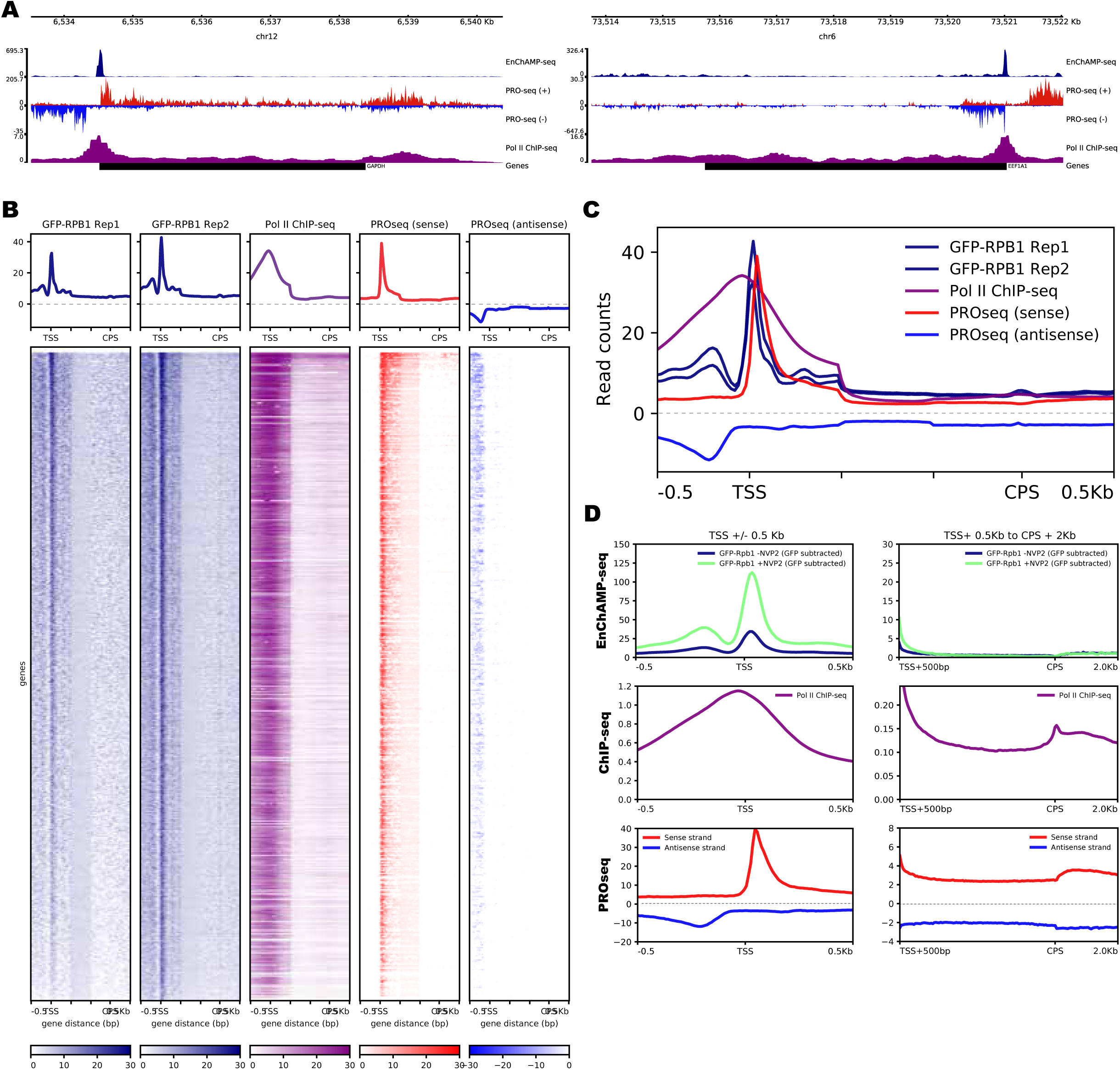
EnChAMP-seq analysis of genomic location/transcription cycle state of EnChAMP captured complexes. A) Genome browser snapshots of two representative genes, GAPDH (left panel) and EEF1A1 (right panel), showing EnChAMP-seq (navy, top track), PROseq (red – plus strand and blue – minus strand, middle track), and Pol II ChIP-seq (purple, bottom track). Gene boundaries from TSS to CPS are shown as a black line at the bottom. GAPDH is a plus strand gene, transcribed left-to-right, and EEF1A1 is a minus strande gene, transcribed right-to-left. Genomic coordinates are shown on top. B) Heatmap plots of two GFP-RPB1 EnChAMP-seq replicates (navy, two left heatmaps), Pol II ChIP-seq (purple), and PROseq (red) for all expressed genes in HCT116 cells. Tick marks represent +/- 500bp window from TSS and CPS, and these regions plotted unscaled. Genebody (TSS+500bp to CPS-500bp) is scaled down to a 500bp window in the middle, for representation purposes, leading to distortion in the signal for all assays. For PROseq, only the sense strand data is shown for simplicity. Average profiles are plotted on top of each heatmap. C) Overlayed profile plots of EnChAMP-seq replicates (navy), Pol II ChIP-seq (purple), and PROseq (red) for all expressed genes as plotted in B from TSS-500bp to CPS+500bp. D) Metagene profile plots of EnChAMP-seq (top panels), Pol II ChIP-seq (middle panels), and PROseq (bottom panels). GFP-RPB1 -NVP2 (navy) and GFP-RPB1 +NVP2 (green) EnChAMP-seq data were plotted after subtraction of the corresponding GFP control sample EnChAMP-seq data. Left panels show the TSS +/- 500bp region, and right panels show the TSS+0.5Kb to CPS+2Kb region where the TSS+0.5Kb-to-CPS region is scaled to 5Kb.

EnChAMP-seq shows a high degree of correlation between replicates (**Fig. 3B**). Although the overall signal patterns of EnChAMP-seq, ChIP-seq, and PROseq look similar –dominated by the TSS-proximal signal –, there are notable differences especially between EnChAMP-seq and ChIP-seq (**Fig. 3B and 3C**). EnChAMP-seq, like PRO-seq, captures both the sense strand Pol II and the upstream divergent Pol II, while the ChIP-seq method lacks the resolution to distinguish these two Pol II signals. Compared to PROseq, the EnChAMP-seq peak is found slightly closer to TSS, with appreciable signal upstream of the TSS. This shift is not caused by the upstream divergent transcription since EnChAMP-seq readily captures that as a distinct peak 200bp upstream of the gene TSS (**Fig. 3C**). It should be noted that while PRO-seq can only detect transcriptionally engaged Pol IIs, both ChIP-seq and EnChAMP-seq can capture chromatin-bound Pol IIs.

Treatment with NVP2 (1.5μM), a selective CDK9 inhibitor, for 1 hour resulted in the accumulation of RNA Pol II at promoter-proximal regions, consistent with the inhibition of pause release^67–69^. EnChAMP-seq analysis indicated that while Pol II signal remained stable and increased at promoters, no signal or noise indicating a decrease in gene body occupancy was detected (**Fig. 3D**). ChIP-seq and PRO-seq show clear signals in the genebody where EnChAMP-seq signal is negligible (**Fig. 3D**). The small increase in early genebody signal observed upon NVP-2 treatment is likely caused by slow dribbling of accumulated paused Pol II with improper post-translational modification (i.e., phosphorylation) after P-TEFb inhibition.

Taken together, the detection of PIC and pause components and lack of transcription elongation factors in GFP-RPB1 EnChAMP-MS (**Fig. 1D and 1F**), lack of any difference in Pol II interactions upon NVP2 treatment (**Fig. 2B**), and lack of genebody signal change upon NVP2 treatment in GFP-RPB1 EnChAMP-seq (**Fig. 3D**), strongly suggests that purification of GFP-RPB1 by EnChAMP predominantly captures factors associated with Pol II in PIC and paused states, the two major rate-limiting steps in the transcription cycle, and thus enabling a deeper study of these steps. The existing structures of mammalian RNA Pol II, representative of various stages in the transcription cycle, provide a plausible explanation why elongating Pol II may not be captured by GFP-RPB1 EnChAMP (**Supp.** Fig. 6). When modelled into the Pol II structures, the GFP-tag on N-terminus of RPB1 is readily accessible in both PIC and paused Pol II structures. However, a well-known elongation factor SUPT6H/SPT6, when modelled as a full-length protein or its partially resolved structures from cryo-EM data, would overlap with GFP-tag on RPB1 N-terminus, making the GFP-tag inaccessible to nanobodies used for purification in EnChAMP.

### EnChAMP-seq provides footprints of RNA Pol II at PIC and Pause states

Our EnChAMP-seq method resembles ChIP-seq in that both identify chromatin fragments bound by target proteins/complexes; however, there are notable differences. Unlike ChIP-seq, EnChAMP-seq does not require cross-linking, and purifications are done under near-native conditions. Benzonase digestion of chromatin yields finer fragmentation of chromatin while preserving weak interactions, with ∼100 bp fragments on average compared to ∼250 bp by sonication in ChIP. In EnChAMP, only DNA fragments bound by the protein of interest are protected from benzonase digestion. Proteins and other large molecular-weight complexes that associate with the target can also contribute to the benzonase protection, thus yielding a “footprint” of the whole complex.

Observing an EnChAMP-seq signal in GFP-RPB1 cells upstream of TSS intrigued us. Upon closer inspection of the actual chromatin fragments mapped around TSS of highly expressed genes, i.e., GAPDH (**Fig. 4A**), we observed two major classes of reads. First class mapped from TSS-50bp to TSS, and second class was mapping from TSS to TSS+50bp. These reads are consistent with expected location of PIC and paused Pol II, respectively. There were some reads that even traversed from TSS-30bp to TSS+50bp suggesting either GTFs associated with Pol II in PIC extending downstream of TSS and providing protection to the downstream DNA, or the elongating Pol II maintaining contact with GTFs upstream of TSS and providing protection. Inspection of chromatin fragments obtained from NVP2 and Triptolide treated GFP-RPB1 EnChAMP samples further supported the assignment of these major read classes to PIC and paused Pol II (**Supp.** Fig. 5); NVP2 treatment led to an increase in both pause- and PIC-associated reads while Triptolide led to near complete elimination of pause reads while maintaining PIC associated reads.

**Figure 4.**
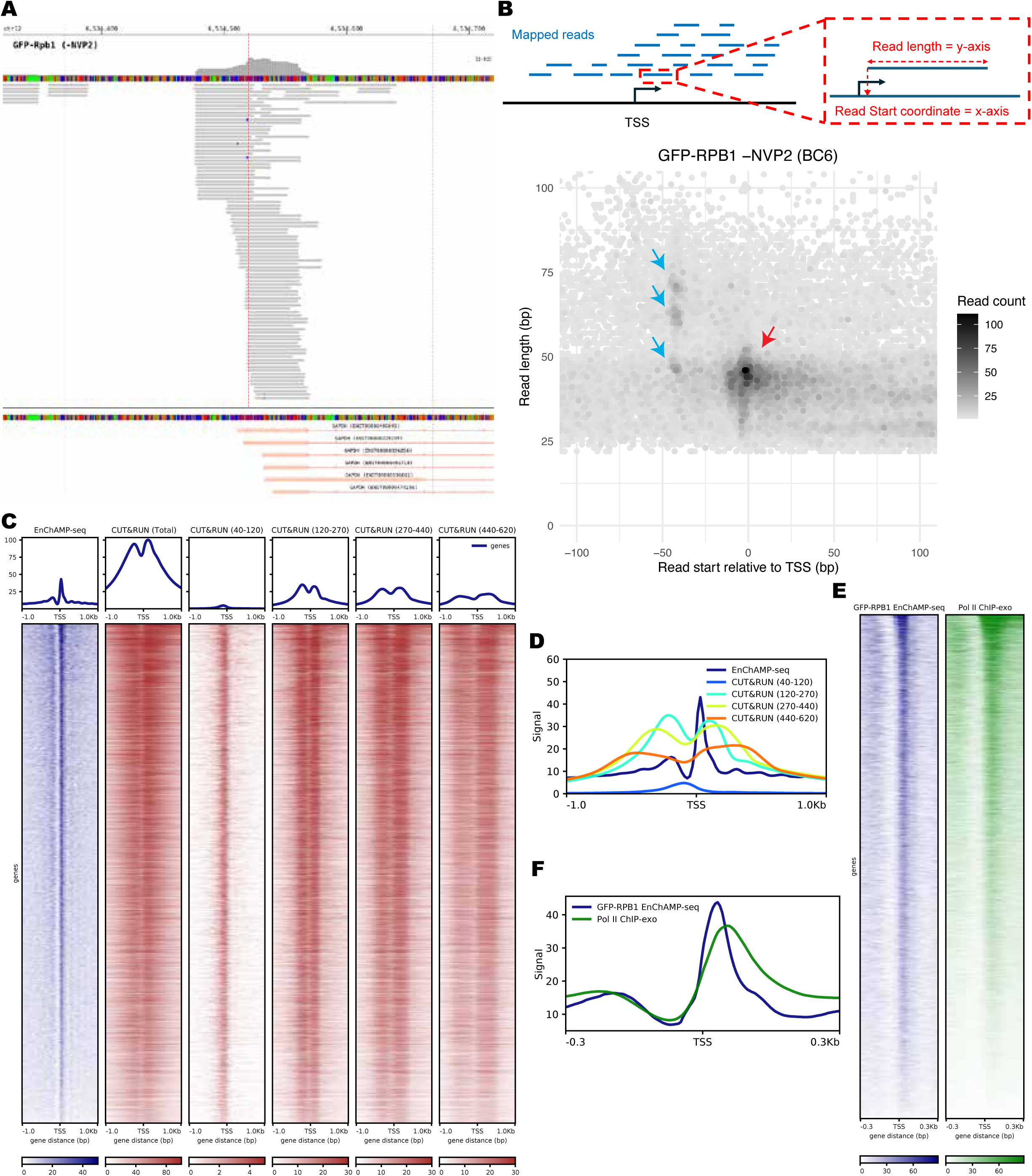
EnChAMP-seq analysis of DNA footprint of RNA Pol II. A) Genome browser snapshot of reads mapped near the TSS of GAPDH gene. The red dashed line marks the PROcap detected major TSS of GAPDH gene, whereas the gray dashed lines mark the +/-150bp from TSS. B) Analysis of DNA footprint of RNA Pol II from GFP-RPB1 EnChAMP-seq data. The top schematic shows how mapped reads were analyzed to visualize the footprint. Read start coordinate relative to PROcap determined gene TSS is plotted on x-axis, while the read length is plotted on the y-axis. Cumulative readcounts from all HCT116 expressed genes were plotted in the bottom graph with the aforementioned coordinates. Gray color scale is used to indicate cumulative readcounts at each position. Blue arrows mark the PIC footprints, while the red arrows mark the paused Pol II footprint. C) Heatmap plots of GFP-RPB1 EnChAMP-seq (navy, left heatmap), and A549 cell Pol II CUT&RUN (brown, right five heatmaps) all expressed genes. CUT&RUN data were plotted either as a whole or separated into different read length classes: Total, 40-120bp, 120-270bp, 270-440bp, and 440-620bp. TSS+/-1Kb region is shown. Average metagene profiles are plotted on top of each heatmap. D) Overlayed metagene profile plots of EnChAMP-seq (navy) and size separated A549 Pol II CUT&RUN data (blue – 40-120bp, teal – 120-270bp, yellow-green – 270-440bp, and orange – 440-620bp) for all expressed genes as plotted in C for TSS+/-1Kb region. E) Heatmap plots of GFP-RPB1 EnChAMP-seq (navy, left heatmap), and K562 cell Pol II ChIP-exo (green, right heatmap) of all expressed genes for the TSS+/-300bp region. Average metagene profiles are plotted on top of each heatmap. F) Overlayed metagene profile plots of EnChAMP-seq (navy) and K562 Pol II ChIP-exo data (green) for all expressed genes as plotted in C for TSS+/-300bp region.

To analyze the potential footprints of Pol II together with the associated complexes genome-wide, we counted and plotted all the reads that were mapped near TSS of expressed genes with the mapped read’s start position relative to TSS on the x-axis and the read-lengths on the y-axis (**Fig. 4B**). This plot revealed a major accumulation of signal for reads starting at TSS with an average length of ∼45bp, suggestive of paused Pol II footprint and consistent with past Pol II footprinting experiments showing ∼50bp protection by Pol II^70–73^. Interestingly, three distinct peaks were observed upstream of TSS, all with starting coordinates −42±2bp relative to TSS, but with varying footprint lengths: 47±2, 62±2, and 71±2bp. Upstream position of these peaks are consistent with footprints of PIC, and suggests that PIC may transition between 3 major configurations during transcription initiation. Similar footprinting analysis of EnChAMP-seq data with drug treatments and that of NELFA-GFP and SPT4-GFP cell lines support the assignment of these footprints to PIC and paused Pol II (**Supp.** Fig. 6A-F). NVP2 treatment does not affect PIC footprint, while increasing the paused Pol II footprint (**Supp.** Fig. 6A and 6B). NVP2 treatment leads to a discernable, but slightly more diffuse, peaks starting around TSS+15±5bp with a footprint of ∼45bp, which is consistent with dribbling of some Pol II downstream of the pause^74^. Triptolide treatment simply wipes out the paused Pol II footprint, while maintaining PIC footprint (**Supp.** Fig. 6C and 6D). We note that, between different experiments we observed some variation between the PIC footprints – discrete peaks vs. a smear ranging from 30 to 70bp footprint. It is unclear whether this reflects the dynamic nature of the PIC complex or simply an experimental variation remains to be studied further. Finally, NELFA-GFP EnChAMP-seq data present a footprint of paused Pol II with no observable PIC footprints, consistent with their interaction with transcriptionally engaged Pol II past beyond initiation.

### Comparison of EnChAMP-seq to CUT&RUN and ChIP-exo

CUT&RUN and ChIP-exo assays were reported to provide RNA Pol II footprints^75,76^. CUT&RUN is performed with non-crosslinked nuclei under native conditions, where the target protein is labeled with a specific antibody, and the target bound chromatin fragment is released by digestion with a Protein A-MNase fusion protein^77^. After sequencing, the mapped fragments are grouped into different size classes and the smallest fragment size bin (40-120bp fragments) was reported to reveal target protein footprint. In contrast, ChIP-exo involves chromatin immunoprecipitation following formaldehyde crosslinking, much like ChIP-seq, but the footprints of the target binding on the chromatin fragment is obtained by lambda exonuclease digestion of the chromatin fragment up to the target protein-DNA crosslinks. ChIP-exo has been demonstrated to have much higher resolution than ChIP-seq and accurately predict target protein binding sites in the genome^78,79^.

Due to lack of publicly available Pol II CUT&RUN and ChIP-exo data in HCT116 cells, we compared our HCT116 GFP-RPB1 EnChAMP-seq data against Pol II CUT&RUN data from A549 cells^75^ and Pol II ChIP-exo data from K562 cells^76^. Albeit originating from different cell lines, we observed a good overall correlation between our HCT116 EnChAMP-seq data and A549 Pol II CUT&RUN data (**Fig. 4C**). Total or larger size CUT&RUN fragments (>120 bp) show two peaks flanking the TSS consistent with divergent transcription patterns observed in humans. However, the smaller fragments (40-120bp) show a single peak claimed to show Pol II footprint^75^. Closer inspection of this signal revealed that this single peak is located upstream of the TSS (**Fig. 4D**) between the two peaks observed in EnChAMP-seq. Thus, it is likely to be originating from a nucleosome free region between the two divergent promoters rather than the footprint of the Pol II. Similarly, we observed good correlation between the HCT116 EnChAMP-seq signal and K562 Pol II ChIP-exo signal (**Fig. 4E**). When compared to this publicly available Pol II ChIP-exo data, it seems EnChAMP-seq provides a higher resolution Pol II footprint (**Fig. 4F**).

## Discussion

Due to its fundamental role in every aspect of biology, transcription by Pol II and its regulation by associated factors has been a focal point of numerous genetic, biochemical, genomic, and structural studies. In the last few decades, a wealth of information has been gathered, and many proteins involved in transcription have been discovered and studied in depth. Comprehensive discovery of all Pol II interactors/regulators is a formidable task; however, such efforts have been attempted several times in the past^30,31,80^ but have been clearly incomplete. To the best of our knowledge, EnChAMP is here shown to be the most effective method to date for capturing RNA Pol II-associated proteins within their native chromatin environment. By maintaining endogenous expression levels and preserving chromatin-bound complexes, EnChAMP provides a more comprehensive and physiologically relevant view of the RNA Pol II interactome (**Fig. 1F** and **Supp. Table I)** compared to other studies^30,31,80^. Our EnChAMP method has proven effective in capturing a large battery of Pol II- and NELF-interacting factors and in footprinting these complexes by sequencing from the highly regulated steps of transcription initiation and promoter-proximal pausing. The identified interactors include both the well-known players and many factors whose functions in transcription regulation are poorly understood. Our study provides a rich source of hypothesis generating data that is useful for the broader transcription community and a technology to study associations of other Pol II interactors.

The keys for EnChAMP success are 1) endogenous tagging of native proteins, thus keeping the expression levels of our bait proteins native and the stoichiometry of all its complexes intact; 2) a carefully-optimized chromatin-isolation protocol to only purify proteins and complexes actively functioning on the chromatin, which is especially important because ∼50% of Pol II molecules in the cell are not bound on chromatin^29^; 3) our EnChAMP purification is done under native conditions, and, unlike other methods, no crosslinking or harsh conditions are involved. ChIP, while ostensibly a competitive alternative method, is not an efficient way to find other proteins associated with bait proteins using proteomics, because in ChIP, target protein is immunoprecipitated from whole cell lysate, under partially-denaturing conditions. This is confirmed by comparing our EnChAMP-MS results of GFP-RPB1 purifications with other published studies using whole cell lysates^30,31^.

In its current form, EnChAMP samples appear to be too heterogeneous and/or at insufficient quantities for structural studies by Cryo-EM. We believe EnChAMP can be readily scaled up and further optimized by sequential purification with dual tagging strategy to enable purification of target complexes at sufficient quantity and purity for structural studies. Recently two groups have successfully purified Pol II from human cells and Drosophila under native conditions and determined Cryo-EM structures from these native complexes^81,82^. Both studies revealed EC structures that include Pol II-Nucleosome-DNA, but the structures lack elongation factors such as SPT6 and PAF complex. Both structures lack direct Pol II-nucleosome interactions providing an explanation for why we did not detect histones in EnChAMP (**Fig. 1F**).

### EnChAMP captures Pol II in PIC and Pause

GFP-RPB1 EnChAMP predominantly captures Pol II in PIC and pause states (**Fig. 1, 2, and 3**). PIC assembly/initiation and promoter-proximal pausing are the two-major rate-limiting steps in transcription cycle, therefore this biased capture of Pol II is a welcomed feature of the current EnChAMP method enabling us to analyze these two steps in greater detail with higher sensitivity. However, EnChAMP appears to be less effective in capturing Pol II and associated factors during elongation. Elongating Pol IIs have to overcome many barriers including nucleosomes, and a number of factors that aid Pol II in overcoming these barriers. PAF complex, a well-established Pol II elongation factor, was not identified as a significant Pol II interactor (**Fig. 1F** and **Supp. Table II**). The lack of elongation complexes in EnChAMP-MS and genebody signal in EnChAMP-seq could be due to several reasons (or various combinations); 1) structural hindrance of the GFP epitope in elongating Pol II complexes (EC), 2) relatively low density of ECs compared to Pol II in PIC and pause states (signal-to-noise ratio), 3) the transient nature of elongation factors interactions with elongating Pol II, 4) a possible sensitivity of elongation complexes to benzonase treatment, and 5) the shear number of different elongation factors utilized in different regions/genes could be diluting their signal in EnChAMP-MS. These speculations need to be further studied. Future tagging of other subunits of Pol II or other factors implicated in other stages of transcription (i.e., elongation or termination) alone or in combination might enable study of these transcriptional stages by EnChAMP. In a recent study, the Adelman endogenously tagged Spt5, DSIF subunit, and performed IP-MS^83^. However, they did not detect many more elongation factors than what is described here for GFP-RPB1, perhaps indicative of the instability of these interactions as has been shown for the PAF1 complex^61^.

### EnChAMP identifies many Pol II interactors whose function in transcription is not well-understood

EnChAMP-MS identified many well-known Pol II interactors including; GTFs, DSIF, NELF, Mediator, and Integrator complexes, building confidence in our purifications and the relevance of the other factors in transcription and its regulation. Our list of high-confidence RPB1 interactors also includes ∼20 proteins involved in rescuing stalled and arrested RNA Pol II elongation complexes that have encountered various impediments during early elongation and in effect have created roadblocks: TFIIS (Tcea1 or Tcea2), Integrator complex, ARMC5, PCF11, RECQL5, and WWP2. The Integrator, a multisubunit complex that interacts with paused Pol II, can terminate early stalled transcription complexes through the activity of an RNA endonuclease subunit that cleaves Pol II’s nascent RNA. The unprotected non-capped 5’ end provides an entry point for exonucleases such that destabilize the elongation complex^41,84,85^. The ARMC5-Cul3 complex acts in a manner complementary to Integrator by ubiquitylation of stalled Pol II targeting it for proteasomal degradation^50,51^. PCF11, a component of cleavage and polyadenylation complex, has been shown to play a role in Pol II termination^86^. Recql5 is a DNA helicase and the only member of the human RecQ helicase family that directly binds Pol II; moreover, it allosterically induces Pol II towards a post-translocation state and may help restart Pol II elongation^87^. WWP2 is a HECT E3 ubiquitin ligase that targets various transcriptional regulators^88^ and has been shown to remove Pol II from double-strand breaks in expressed genes to aid DNA repair by preventing collision between DNA repair and transcriptional machineries^89,90^. Thus, a rich and diverse battery of mechanisms exist to resolve Pol II stalling or arrest, which can arise when Pol II transcribes normally and is particularly pronounced when Pol II confronts various obstacles in its path^91^.

ARMC5 was the subject of a pair of recent elegant studies showing it functions like Integrator to terminate RNAs in the early elongation to pausing stages^50,51^. These two labs have made impressive progress recently on the roles of Integrator and the ARMC5 directed CUL3 ubiquitin ligase showing they are likely acting redundantly to remove Pol II that is in peril, either irreversibly stalled or in co-directional collision with replication machinery^92^. Our Rpb1 pull-downs following treatment of cells with the TFIIH helicase inhibitor triptolide, which blocks Pol II during early phases of initiation, robustly recruits the CUL3 ubiquitin ligase^50^ and the proteasome (**Fig. 2D**), but not the WWP2 ubiquitin ligase found reproducibly to associate with RPB1 in normally grown cells. The ARMC5-CUL3 recruitment likely accounts for known degradation of Pol II during triptolide inhibition^93^. All these mechanisms of removing a Pol II blockade on the DNA template are likely coordinated but in ways that are far from fully understood. Future studies that address features of their interplay at the transcription, chromatin, and factor binding levels are needed.

The PAQosome is a 12-subunit chaperone complex^48^ involved in assembly of various protein complexes including Pol II^49^. Interestingly, Rpb5/Polr2E, a subunit of RNA Pol II, is considered a component of PAQosome complex. The entire collection of 12 subunits that form the PAQosome chaperone are reproducibly identified as RPB1 interactors in our MS analysis (**Supp. Table II**). These include the RUVBL1 and RUVBL2 ATPases, which also function in other complexes^94^, that drive conformational changes in client proteins, the RPAP3 and PIH1D1 that connect to various substrates such as RNA Pol II^95^. We speculate the driving conformational and compositional changes allow the transition of RNA Pol II complexes from one step in the transcription cycle to the next, perhaps analogous to those transitions seen in the helicase-driven steps of the splicing cycle.

Interestingly, we also find several other factors that are known to have a role in Pol II assembly and transport to the nucleus, but also are reproducibly found associated with RPB1 on chromatin. These include GPN3 and RPAP2, whose cryo-EM structure with Pol II has recently been determined, suggesting that it creates a checkpoint for initiation^42,43^. RPAP2 is also associated with an RPRD-associated S5 phosphatase complex that acts on the CTD of Rpb1^44^. We propose that a large battery of chaperones interacting with chromatin bound RPB1 might participate in the assembly and disassembly of the multiple complexes with different compositions that are required for Pol II to progress through the transcription cycle.

The mechanistic roles of many factors that have been enriched by GFP-RPB1 or GFP-NELFA EnChAMP are unknown. These include several zinc binding proteins (Zmynd8, Znf592, Znf609, Znf655, and Znf687) that have been implicated in diseases and transcription^96–98^, but their roles are as yet far from being understood. They all are detected reproducibly in GFP-RPB1 pull-downs and all but Znf687 are detected in NELFA pull-downs as well. Thus, this set of zinc-binding proteins warrant investigation based on their associations with disease, with promoter-proximal pausing, and because their mechanism of action in transcription is not understood.

Based on our EnChAMP-seq results, we suspect most if not all of the identified LMCs interact with Pol II in the PIC or at the pausing stage. It is highly unlikely that all of these LMCs interact with Pol II at the same time, or even at the same genomic loci. Future studies of individually tagged LMCs by EnChAMP-MS and -seq assays will identify the composition of these LMCs and the other LMCs that co-associate with Pol II, and determine where on the genome, which genes and what stage of the transcription cycle, that they interact with Pol II. Thus, these LMC-targeted EnChAMP analyses will assess the stage of transcriptional cycle and the genomic locations where Pol II interactions occur, and thereby also determine whether different LMCs interact with Pol II at the same or different times, and at the same or distinct genomic loci.

Interestingly, some of the EnChAMP-MS identified Pol II interactors have been implicated to function in other steps of the transcription cycle in addition to their well-established functions. For example, the FACT complex, generally considered an elongation factor, has recently been shown to play a role in Pol II pause release^99,100^; and PCF11, component of the cleavage and polyadenylation complex, might also function at the pause region^86,101,102^. These seemingly misplaced interactions suggest two interesting possibilities: either i) these factors are pre-loaded onto the transcriptional machinery much earlier than their function is needed, or ii) these factors have additional functions that are underappreciated at different stages of the transcription cycle. A detailed study of these factors with EnChAMP and other complementary assays is warranted to elucidate their transcription regulatory mechanism.

Pol II CTD is known to undergo post-translational modifications by a number of kinases during the transcription cycle ensuring timely and coordinated transcriptional responses^10,103^. Ser5 phosphorylation by the CDK7 subunit of TFIIH complex during initiation and Ser2 phosphorylation by P-TEFb are well studied and serve as a start signal for transcription initiation and pause release, respectively. CDK9 must phosphorylate Pol II and associated factors to release Pol II from promoter-proximal pausing to productive elongation. Upon termination, phosphatases remove these phosphorylation marks to enable recycling of the transcription complex components for re-initiation at other genes^104^. It is less well understood how transcriptional kinases and phosphatases regulate processive elongation by Pol II. The kinases CDK12/13 are thought to phosphorylate Pol II during elongation^105^, and CDK9 may directly bind elongating Pol II via the super-elongation complex^106^. Our EnCHAMP-MS data indicate that multiple phosphatases—RPAP2, CTDP1, and PP2A—interact with paused Pol II, suggesting that pause release might be regulated in a more subtle manner than previously appreciated by the opposing actions of kinases including CDK9, and phosphatases. However, we know little about how ongoing phosphorylation and dephosphorylation regulate elongation—specifically whether continued phosphorylation of Pol II underlies the gradual speedup of transcription during the first several kilobases of the elongation phase^12^.

### GFP-NELFA EnChAMP-MS Reveals Protein Interactions at RNA Polymerase II Pausing Sites

Using GFP-NELFA EnChAMP-MS, we identified a targeted set of 83 proteins associated with RNA Pol II pausing (**Fig. 2C** and **Supp. Table II**). Among these proteins, the Negative Elongation Factor (NELF) complex, comprising NELFA, NELFB, NELFC, and NELFD, emerged as central in stabilizing RNA Pol II at gene promoters to prevent premature transcription elongation^5,107,108^. Additionally, cofactors SPT4 and SPT5, components of the DRB Sensitivity-Inducing Factor (DSIF), were enriched, highlighting their direct roles in regulating transcriptional pausing^109,110^. Several Integrator complex proteins were enriched as expected, since the Integrator has a role in RNA cleavage and termination of paused Pol II at some genes^111,112^. The identification of additional factors, including CMTR1 and PCIF1, suggests a close interplay between RNA pausing, mRNA capping, and subsequent processing events^113–115^.

Compared to GFP-RBP1 EnChAMP-MS, which detected 184 interacting proteins, GFP-NELFA revealed a more selective set of interactors at pause, reflecting functional specialization at the pausing phase of the transcription cycle. NELFA-GFP EnChAMP-seq reveals a footprint consistent with paused Pol II, without any footprint at PIC. This distinction is critical in understanding why Mediator does not exhibit the expected enrichment following Triptolide treatment, despite its role as a key component of the PIC^17^. If Mediator were stably associated with the stalled PIC, an increase in its signal would be expected after Triptolide treatment, similar to other PIC-associated factors such as TFIIH and TFIID^116^. However, our data do not show this enrichment. Instead, Mediator levels remain unchanged in GFP-RPB1 immunoprecipitation, suggesting that Triptolide treatment alters the PIC in a way that weakens or disrupts Mediator association. Recent structural studies have demonstrated that Mediator directly interacts with the unphosphorylated CTD of RNA Pol II within the preinitiation complex, playing a crucial role in stabilizing early transcription assemblies^17^. A likely explanation for our observation is that Triptolide extends the lifespan of the stalled PIC, allowing additional phosphorylation events on the RPB1 CTD that destabilize Mediator binding. TFIIH, which remains active even when XPB helicase is inhibited, contains CDK7—a kinase known to phosphorylate Ser5 of the CTD repeats^117,118^. If the stalled complex persists long enough, it may provide an opportunity for further CTD phosphorylation, potentially by CDK9, reinforcing Mediator dissociation before elongation begins^119^. Taken together, these results highlight the compositional and positional transformations that take place as Pol II progresses from one regulated state to the next, with NELF complex function restricted to the pause state.

### Comparative Analysis of RPB1 Interactome Changes Mediated by Triptolide and NVP2

The transcriptional consequences of Triptolide (inhibition of transcription initiation) and NVP2 (inhibition of pause release) reveal distinct mechanisms in RNA polymerase II regulation. Triptolide treatment leads to a comprehensive reduction in RNA Pol II occupancy at promoters, consistent with its role in inhibiting transcription initiation^120^. In contrast, NVP2 enhances promoter-proximal pausing without diminishing Pol II, consistent with its role in inhibiting P-TEFb phosphorylation of the paused Pol II complex and thereby blocking release of Pol II to productive elongation without interfering with initiation^59^. These findings delineate the sequential nature of transcription initiation and elongation control mechanisms, underscoring the value of chemical perturbation approaches in studying Pol II transcriptional dynamics.

Despite its dramatic effect on transcription, NVP2 treatment had no significant effect on Pol II interactome as identified by GFP-RPB1 EnChAMP-MS. While the EnChAMP-seq assessed Pol II footprint signal at both PIC and pause were increased. These results are consistent with a predominant capture of Pol II at PIC and pause by our EnChAMP method. Triptolide treatment on the other hand induced a pronounced shift in the RPB1 interactome, significantly impacting transcriptional regulation and protein stability. Many PIC-associated factors i.e., TAF5, TAF7, TAF8, and TAF9 maintain their association with RPB1 even after triptolide treatment, indicating the persistence of certain complex elements despite initiation inhibition. However, Triptolide disrupts interactions with critical factors involved in transcription elongation and RNA processing, such as PHF8, CMTR1, SSRP1 and SUPT5H. EnChAMP-Seq analysis validated findings from EnChAMP-MS, demonstrating a complete elimination of RNA Pol II occupancy at promoter proximal pause region, while maintaining or increasing occupancy at PIC following Triptolide treatment.

Additionally, Triptolide treatment redirects the RPB1 interactome toward a degradation-associated profile, characterized by the enrichment of proteasomal and ubiquitin-related proteins, including ADRM1, PSMD1, PSMD3, PSMC2, and CUL3. Many of these proteins are components of the 26S proteasome and ubiquitin-proteasome system, indicating a cellular response targeting stalled Pol II for degradation. ADRM1, a proteasomal ubiquitin receptor, regulates protein turnover and homeostasis, while ARMC5 participates in transcriptional regulation and cellular stress responses. BRD2, belonging to the bromodomain and extra-terminal (BET) family, facilitates chromatin remodeling and transcriptional activation by recognizing acetylated histones (**Table 2**). These findings collectively suggest that Triptolide induces RPB1 degradation via the ubiquitin-proteasome system, indicating a quality control mechanism that eliminates stalled polymerases to maintain transcriptional homeostasis.

### Why some well-known transcription-associated factors are missing in EnChAMP-MS

Among the EnChAMP-MS identified high confidence Pol II interactors, there are some notable missing factors (**Fig. 1F** and **Supp. Table X**). P-TEFb, master regulator of pause release on at least 95% of the expressed genes^12^, was not detected in any of EnChAMP-MS experiments including the drug treatments as a significant Pol II or NELF interactor. TFIIB, which functions as a bridge between TBP and Pol II and positions Pol II for proper transcription initiation, although detected, it was not enriched as a Pol II interactor in EnChAMP-MS experiments except upon Triptolide treatment. TATA Binding Protein (TBP), a critical component of TFIID complex and responsible for DNA bending and assembly of PIC, was not readily identified as a Pol II interactor. Additionally, whole complex (i.e., PAF complex) or individual subunits of various complexes (i.e., CDK8 subunit of Mediator and SUPT16H/SPT16 subunit of FACT complex) are also missing among the EnChAMP-MS identified Pol II interactors. Some of these are due to not meeting our stringent enrichment criteria, while others are not detected as a significant Pol II interactor even in a single EnChAMP experiment. Note that the GFP-RPB1 and NELFA-GFP EnChAMP-MS experiments were repeated 3 times each with 4 biological replicates against the GFP control samples for the untreated samples. For example, SPT6, an elongation factor that interacts with Pol II and/or chromatin tightly^61^, and XRN2, a 5’-to-3’ exoribonuclease that terminates transcribing Pol II following Cpsf73-mediated cleavage^121^, did not meet our stringent criteria (identified in 2 out of 3 GFP-RPB1 EnChAMP-MS experiments). While others are absent among the significant interactors in any of the experiments, likely because of the transient nature of their interaction with Pol II or instability leading to a loss during EnChAMP. For example; both P-TEFb^61^ and PAF1^61^ are known to interact with Pol II transiently^61^.

Interestingly, histone proteins were not identified as significant interactors of either Pol II or NELF complexes by EnChAMP-MS. A number of previous studies implicated nucleosomes as a barrier for Pol II and responsible for its pausing; however, the lack of histone protein enrichment in EnChAMP experiments, the distance between paused Pol II and the +1 nucleosome as measured by PRO-seq or EnChAMP-seq determined Pol II position and the MNase-seq determined nucleosome position, suggest that paused Pol II (or Pol II in the PIC) do not form stable interactions with nucleosomes, If nucleosomes play a major role in establishing the pause, it may be as an elongation energy barrier rather than through specific interactions. Recent Cryo-EM studies of Pol II purified from native sources determined structures of elongating Pol II-Nucleosome complex; however, the interaction of Pol II with the nucleosome is minimal. It remains to be seen if these structures represent transient semi-stable tripartite structures formed by elongation factors/histone remodelers-Pol II-nucleosome during elongation in which the elongation factors/histone remodelers were lost during purifications.

### Footprinting RNA Pol II with EnChAMP-seq

EnChAMP-seq provides a high resolution genome-wide footprint of RNA Pol II in PIC and pause states. Previous DNase I-based studies have shown a ∼50bp footprint for paused Pol II, which is consistent with EnChAMP-seq derived ∼45bp footprint^73,122^. NVP2, a Cdk9 kinase inhibitor, is known to increase pause and cause some dribbling of paused Pol II downstream of its normal pause position. Our analysis indicates that the footprint of these two Pol IIs are very similar in size, suggesting minimal structural change in the extent of DNA interactions between these two types of Pol IIs. More interestingly, EnChAMP-seq was able to footprint Pol II in PIC, to the best of our knowledge for the first time, and revealed a distinct set of footprints. In some experiments, we detected three distinct footprints while in others a more of a continuum ranging from 30bp to 70bp footprints. This suggests that during PIC assembly and transcription initiation Pol II undergoes dynamic structural changes with some semi-stable transitional states.

In summary, our EnChAMP-MS method provides to date the largest set of Pol II interactors at PIC and pause, the two major rate-limiting steps of the transcription cycle. EnChAMP-seq complements this with high-resolution footprinting of Pol II. Many of the identified Pol II interactors are understudied and suggest intricate regulation of Pol II transcription via multiple mechanisms. Therefore, this data and our method will foster future studies of how these factors interplay to regulate transcription.

## Materials and Methods

### Generation of Endogenously Tagged Cell Lines

All endogenously tagged cell lines (GFP control, GFP-RPB1, and NELFA-GFP) used in this study were generated by CRISPR/Cas9 gene editing of the HCT116 parental diploid cell line^123,124^. Briefly, gene specific guide RNA (gRNA) sequences were cloned into pX330 plasmid^125^, while the Homology Directed Repair (HDR) templates targeting each gene were constructed in pUC19 plasmid. HDR constructs were created by flanking the insertion cassettes with up to 1Kb genomic fragments of either side of start or stop codons for N- and C-terminal tagging. RPB1 is N-terminally tagged with a GFP insert. GFP control cells were generated by inserting a GFP-P2A sequence upstream of the RPB1 coding sequence. NELF-A was C-terminally tagged with a GFP insert followed by a PGK promoter driven Blasticidin Resistance gene, cloned from pMDD54_EGFP_Bsr_V2 plasmid, Supplementary Material). The left and right homology arm (LHA or RHA), up to 1Kb fragments, were PCR amplified from genomic DNA obtained from HCT116 cells. Assembly of LHA-GFP insert-RHA in the pUC19 backbone was achieved either by restriction enzyme digestion/T4 DNA ligase reaction or by Gibson Assembly^126^. All gRNA sequences and oligos used for construction of HDR constructs and genotyping the resulting cell lines are listed in **Supplementary Table I**, and complete sequences of HDR constructs are included in **Supplementary Materials**.

Parental and engineered HCT116 cells were cultured in McCoy’s 5A media (Thermo Fisher Scientific, Cat# 16600-108) supplemented with 10% Fetal Bovine Serum (VWR, Cat# MP1300500H) and 100 U/ml Pen-Strep (Thermo Fisher Scientific, Cat# 15140122) in a humidified cell culture incubator at 37 °C with 5% CO_2_ atmosphere.

To generate tagged cell lines, gene specific gRNA expression construct and the HDR template constructs (1:4 ratio) were co-transfected using FuGene HD reagent (Promega, Cat# E2311) at DNA:FuGENE HD reagent ratio of 1:4 following manufacturer’s protocol in 6-well plates. 24h post-transfection cells were split and cultured for an additional week, in the presence of 8 μg/ml Blasticidin (Thermo Fisher Scientific, Cat# R21001) for SPT4 and NELF-A or absence for all others. GFP expressing single cell clones were selected by Fluorescence Activated Cell Sorting (BD Biosciences FACSAria Fusion instrument) at Cornell Institute of Biotechnology Flow Cytometry Facility.

Individual clones were tested by genotyping PCR and/or by Western Blot using target specific antibodies (RPB1: 8WG16, Thermo Fisher Scientific, Cat# MA1-10882, RPB2: ProteinTech, Cat# 20370-1-AP, RPB3: ProteinTech, Cat# 13428-1-AP, NELF-A: ProteinTech, Cat# 10456-1-AP, SPT4: Santa Cruz Biotechnology, Cat# sc-515238), or anti-GFP antibody (Abcam, Cat# Ab290). For each target protein, a single clone that was determined to be homozygous was selected for all subsequent experiments.

### EnChAMP Protocol

#### Cell Culture and Harvesting

HCT116 GFP-RBP1 cells were cultured in 15 cm plates and treated 1.5 μM of NVP-2 (MedchemExpress, Cat# HY-12214A) for 1 hour, 1 µM of Triptolide (Millipore Sigma, Cat# T3652) for 20 minutes, or left untreated. Cells were harvested at ∼90% confluency (∼20 million cells/plate). Cells were washed with DPBS, scraped, and collected by centrifugation at 800g for 5 minutes at 4°C. Supernatants were discarded.

#### Nuclei Isolation

Cell pellets were resuspended in 1 mL of ice-cold Hi-C Buffer A (10 mM HEPES [pH 7.5], 10 mM KCl, 1.5 mM MgCl_2_, 1 mM CaCl_2_, 0.1% Digitonin (Millipore Sigma, Cat# 300410), with 1X Phosphatase (Sigma Aldrich, Cat# P2850) and 1X Protease inhibitor cocktails (Thermo Fisher Scientific, Cat# A32965), and 1 mM PMSF (Cell Signalling, Cat# 8553S)) and incubated on ice for 30 minutes to allow cell swelling. After incubation, aliquots were taken for western blot analysis. The remaining lysates were centrifuged at 1000g for 15 minutes at 4°C. Nuclei pellets were washed with 1 mL of cold Hi-C Wash Buffer B (Hi-C Buffer A supplemented with 250 mM sucrose), centrifuged as above, and supernatants containing cytoplasmic fraction was discarded.

#### Chromatin Isolation

Nuclei pellets were resuspended in 500 μL of 1.5% Digitonin Chromatin Isolation Buffer A (10 mM HEPES [pH 7.5], 20 mM KCl, 150 mM NaCl, 1.5 mM MgCl_2_, 1 mM CaCl_2_, 15% Glycerol, 1.5% Digitonin, with phosphatase and protease inhibitors, and PMSF) and incubated on ice for 30 minutes. Samples were centrifuged at 1000g for 30 minutes at 4°C, and supernatants containing nucleoplasmic fraction were discarded. The pellets were resuspended in 500 μL of fresh Buffer A, and 3 μL of 250 U/μL Benzonase® Nuclease (Millipore Sigma, Cat# E1014) was added for chromatin shearing. The samples were rotated at 4°C for 1 hour, followed by centrifugation at 1000g for 30 minutes at 4°C. Soluble chromatin proteins were collected as supernatants, while the insoluble chromatin fractions were discarded.

#### Affinity purification with anti-GFP nanobody

During Benzonase digestion of chromatin, per sample 10 μL of GFP-Trap® M-270 magnetic particles (ChromoTek, Cat#gtd) were washed three times with Chromatin Isolation Buffer B (10 mM HEPES [pH 7.5], 20 mM KCl, 150 mM NaCl, 1.5 mM MgCl_2_, 1 mM CaCl_2_, 15% Glycerol) and resuspend in 50 μL of Buffer B. Soluble chromatin proteins (∼500 μL) were incubated with 50 μL of pre-washed magnetic particles per sample overnight at 4°C. Beads were washed sequentially with Chromatin Isolation Buffer B: a 5-minute rotation wash, followed by two stationary washes, and a final transfer to a clean tube. Beads were eluted with 200 μL of 100 mM HEPES (pH 7.5) containing 1% SDS at 65°C for 15 minutes. The supernatants (eluted chromatin) were collected for downstream analysis, and 20 μL was reserved for western blot validation.

After initial optimizations, later EnChAMP experiments were carried out with a scaled down version where ∼10 million cells grown in 10 cm dishes were used per sample. ∼200 μg soluble chromatin proteins were incubated with 5 μL of pre-washed magnetic particles slurry per sample overnight at 4°C and buffer volumes were scaled down accordingly. GFP-Trap® M-270 magnetic particles (NanoTag Biotechnologies, Cat# N0310,) were used due to their lower cost and elutions were done with 100 μL of 100 mM HEPES (pH 7.5) containing 1% SDS at 95°C for 8 minutes. These scaled down EnChAMP experiments were indistinguishable from earlier, larger scale experiments.

### Western blot

GFP-RPB1 or NELF-GFP bait proteins were separated using a 4–15% homemade gradient SDS-PAGE gel and subsequently transferred to PVDF membranes using a wet transfer method. The transfer was performed at a constant current of 300 mA for 90 minutes in Transfer Buffer (25 mM Tris, 192 mM Glycine, 20% Methanol). After the transfer, the membranes were blocked with 5% non-fat dry milk prepared in PBST buffer (1× PBS with 0.1% Tween-20) for 1 hour at room temperature to prevent nonspecific binding. The membranes were then incubated overnight at 4°C with primary antibodies: Rpb1 NTD (D8L4Y) Rabbit mAb (Cell Signaling Technology, Cat# 14958) and NELF-B/COBRA1 (D6K9A) Rabbit mAb (Cell Signaling Technology, Cat# 14894). Each antibody was diluted 1:5,000 in PBST containing 5% non-fat dry milk. Following incubation, the membranes were washed three times with PBST (5 minutes each at room temperature) and subsequently incubated with Alexa Fluor 680 donkey anti-rabbit IgG secondary antibody (Invitrogen, Cat# A-21109), diluted 1:5,000 in PBST containing 5% non-fat dry milk for 1 hour at room temperature (RT). After 3 additional 5 minutes washes with PBST at RT, the protein bands were visualized using an Odyssey infrared imaging system (LI-COR Biosciences). For Western Blot validation of cellular fractionation in EnChAMP, Histone H3 Rabbit pAb (Proteintech, Cat# 17168-1-AP), GAPDH Mouse mAb (Proteintech, Cat# 60004-1-Ig), Beta Actin Mouse mAb (Proteintech, Cat# 66009-1-Ig), and Beta Tubulin Rabbit pAb (Proteintech, Cat# 10094-1-AP) were used.

### Mass spectrometry

#### Sample preparation

The eluted proteins were reduced with 10 mM TCEP (Thermo Scientific, Cat# 77720) for 30 minutes at room temperature and alkylated in the dark using 18 mM iodoacetamide (IAA, Cytiva, Cat# RPN6302). Next, samples were precipitated using 550 μL precipitation solution (PPT, 50% acetone, 49.9 % ethanol and 0.1% acetic acid) and incubated overnight at −20 °C. The mixture was centrifuged at 15000 rpm for 10 min at 4 °C, and the protein pellets were washed twice with 550 μL PPT to remove detergent. After centrifugation, the pellets were air-dried over 30 minutes at room temperature. The dried pellets were dissolved with 30 μL 8M urea in 50 mM Tris-HCl, pH 8.0, then urea was diluted by adding 90 μL 50 mM Tris-HCl, pH 8.0, 150 mM Nacl. Finally, proteins were digested overnight using 500 ng Trypsin Gold (Promega, Cat# V5280) at 37 °C. Digested material was acidified by adding 120 μL of 4% formic acid solution.

#### EvoTip Sample Preparation

EvoTips were conditioned with 100% isopropanol for 1 min, washed two times with 50 μl EvoTip Buffer B (Mass Spec Grade Acetonitrile with 0.1% formic acid (FA)) by centrifugation for 60 seconds at 700 g. Washed EvoTips were equilibrated with three times 50 μl EvoTip Buffer A (Mass Spec Grade Water with 0.1% FA) and centrifugation for 60 seconds at 700g. The sample (120 μL) was loaded onto the EvoTip, followed by centrifugation for 60 sec at 700g. The loaded peptides were washed two times with 120 μl EvoTip Buffer A by centrifugation for 60 sec at 700g each. The washed peptides were kept wet by applying 250 μl of EvoTip Buffer A on top of the EvoTip and centrifugation for 30 s at 700g.

#### Reverse Phase Liquid Chromatography

The peptides on the EvoTips were separated on an Evosep One chromatography system using a homemade 8 cm × 150 μm analytical column, packed with 1.5 μm C18 beads. Peptides separated from the stationary phase over 22 min according to the manufacturer standard method 60SPD. Peptides were eluted from the column with solvent A (Mass Spec Grade Water with 0.1% FA) and gradually increasing concentration of solvent B (Mass Spec Grade Acetonitrile with 0.1% FA).

#### Mass spectrometry

All samples were analyzed on a timsTOF HT (Bruker) Q-TOF mass spectrometer coupled to a Evosep LC system. Samples were run using diaPASEF methods, consisting of 12 cycles including a total of 34 mass width windows (25 Da width, from 350 to 1200 Da) with 2 mobility windows each, making a total of 68 windows covering the ion mobility range (1/K0) from 0.64 to 1.37 V s/cm2. These windows were optimized with the Window Editor utility from the instrument control software (timsControl, Bruker) using one DDA-PASEF run acquired from a pool of the analyzed samples. Briefly, this utility loaded the run and represented its ion density in the m/z and ion mobility ranges (i.e. the mobility heatmap), so the dia-PASEF windows coverage could be adjusted to ensure complete coverage, and the window settings calculated. The collision energy was programmed as a function of ion mobility, following a straight line from 20 eV for 1/K0 of 0.6 V s/cm2 to 59 eV for 1/K0 of 1.6 V s/cm2. The TIMS elution voltage was linearly calibrated to obtain 1/K0 ratios using three ions from the ESI-L Tuning Mix (Agilent) (m/z 622, 922, 1222) before each run, using the ‘Automatic calibration’ utility in the control software (timsControl, Bruker).

#### Data Analysis

The Bruker timsTOF HT instrument was used in DIA-NN version 1.8.1 to analyze the diaPASEF runs. In DIA-NN, missed cleavages were set to 0, precursor change range 2-4, and precursor m/z range 349-1500, neural network classifier set to double-pass mode, quantification strategy was set to ‘Robust LC (high precision)’, and MBR option was enabled. MS1 and MS2 accuracy, and retention time window scans, were set to 0 in order to let DIA-NN to perform their automatic inference for the first run in the experiment. All other DIA-NN settings were left default, using RT-dependent cross-run normalization and filtering the output at 1% FDR. The number of threads used by DIA-NN, were 32, as automatically suggested by the software.

The resulting data were analyzed and visualized using Python, R, and Microsoft Excel. For each target protein, enrichment was assessed relative to the GFP control, each with 4 biological replicates. Ratios were calculated by pairing replicates in a defined manner, and the fold change (FC) was determined as the median of all possible ratios associated for a given protein. To assess statistical significance, an Empirical Bayes approach was applied, comparing each protein’s ratios to those of all other proteins in the dataset to generate raw p-values. These p-values were subsequently adjusted using the Benjamini–Hochberg procedure to control the false discovery rate (FDR).

For EnChAMP-mass spectrometry (EnChAMP-MS) experiments, cells were cultured in separate dishes (N = 4 replicates per condition). Analyses were performed independently three times for both GFP–RPB1 and NELFA–GFP samples, and final interactors were defined as those consistently identified in at least two out of three experiments. Protein interactors for each bait were identified by comparison to GFP control cells, using thresholds of FC ≥ 2 and FDR < 0.05. Known contaminants frequently observed in affinity purification (AP)-MS experiments—such as keratins (KRT), small ribosomal subunit proteins (RPS), and large ribosomal subunit proteins (RPL)—were excluded from the final dataset.

To quantify differences in protein abundance between two experimental conditions (+/-NVP2 or +/-Triptolide drug treatments), we used the pipeline described above to calculate FC and FDR. Protein interactors for each bait were identified by comparison to GFP control cells, using thresholds of FC ≥ 1.5 and FDR < 0.1. The analysis then focused on the union of interactors identified in both untreated and drug-treated bait conditions. To assess drug-specific effects on protein interactions, we performed a direct comparison between drug-treated and untreated bait runs. Data were normalized to the intensity of the bait protein in each sample to account for differences in bait expression levels. Results were visualized using a volcano plot, plotting fold changes against adjusted p-values. Known contaminants were removed prior to downstream analysis. Custom analysis scripts are available upon request.

The mass spectrometry data have been deposited to the ProteomeXchange Consortium via the PRIDE partner repository with the dataset identifier PXXXXX.

### EnChAMP-seq

Eluates of ∼5M cell equivalent from EnChAMP experiments were brought to 300ul with 1X TE buffer (10 mM Tris-Cl pH 8.0, 0.1 mM EDTA), extracted once with equal volume Phenol:Chloroform mix pH 8.0 (Thermo Fisher Scientific, Cat# 17909), extracted with equal volume Chloroform, EtOH precipitated (1/10th volume 3M NaAcetate pH 5.2, 2 μl GlycoBlue co-precipitant (Thermo Fisher Scientific, Cat# AM9516), and 3 volumes 100% EtOH, washed with 70% EtOH, and air dried pellet was resuspended in MilliQ-H2O. DNA amount was quantitated by Qubit dsDNA HS Assay (Thermo Fisher Scientific, Cat# Q32851). 2-40 ng DNA was end-repaired in the presence of a 250 μM dNTP mix (each), 1x T4 DNA Ligase Buffer (NEB), 6 units T4 DNA Polymerase (NEB, Cat# M0203L), 20 units T4 PNK (NEB, Cat# M0201L), and 2.5 units Klenow Polymerase (NEB, Cat# M0210L) at RT for 30 min on a Thermomixer at 600 rpm. End-repaired DNA was purified using MinElute Reaction Clean-up Kit (Qiagen, Cat# 28204) and A-tailed in the presence of 1x Buffer #2 (NEB), 200 uM dATP, and 10 units Klenow exo- polymerase (NEB, Cat# M0212L) at 37C for 30 min on a Thermomixer at 600 rpm. A-tailed DNA was purified using MinElute Reaction Clean-up Kit (Qiagen, Cat# 28204) and ligated to Illumina TRUseq adapters in the presence of 12-120 nM TRUseq Index adapter, 1x T4 DNA Ligase buffer (NEB), 1,200 units T4 DNA Ligase (NEB, Cat# M0202L) at 18 °C O/N. TRUseq adapter ligated DNA was purified using MinElute Reaction Clean-up Kit (Qiagen) and subjected to a pilot PCR using P5 and P7 oligos to determine appropriate number of PCR cycles to generate a library. Final PCR reactions were carried 11-15 cycles and purified using MinElute PCR Clean-up Kit (Qiagen, Cat# 28004). After quality check with BioAnalyzer analysis, if deemed necessary the libraries were gel purified (150-350 bp region) using 8% native PAGE to get rid of the adapter dimer (∼130 bp). Equal amounts of 8-12 samples were pooled and sequenced either by 2x150nt on an Illumina NovaSeq 6000 instrument (NovaGene) or by 2x80nt on an Element Biosciences AVITI instrument (EGC Core Facility).

Sequencing data was analyzed with a custom analysis pipeline. Briefly, adapter sequences were removed using cutadapt (ver3.7)^127^ and mapped to the human hg38 genome using bowtie2 –local (ver2.5.1)^128^. The sam file was converted to an indexed bam file with samtools view, sort, and index (ver1.7)^129^. Finally, the indexed bam file was converted to a RPKM normalized bigwig file using DeepTools bamCoverage (ver3.5.2)^130^.

### Footprinting analysis

Mapped reads within 300bp of the PROcap detected TSSs of expressed genes (**Supp. Data II)** were selected from the indexed bam files with samtools view (ver1.7), converted to a bed file with bamToBed (v2.26.0)^131^, intersected with the PROcap corrected gene TSSs with bedtools intersect (v2.26.0). Genomic coordinates were then transformed to TSS-relative coordinates using common bash tools cut, sort, uniq, and etc (ver5.2.37). Text editing tools awk (ver5.2.1) and sed (ver4.9) were used to edit large text files such as bed files of gene lists when necessary. Footprinting plots were generated in R (ver4.4.2) with the ggplot2 package (ver3.5.2)^132^. PROcap and PROseq data were processed as previously published^133,134^ including the bigWig package (ver0.2-9) in R. All R scripts were run in RStudio (2023.03.2 Build 454).

The list of HCT116 cell line expressed genes with the PROcap corrected TSS coordinates in the bed format is provided as a **Supp. Data II**. Top 10% highest HCT116 expressed genes (N=1,300) were selected based on their PROseq derived Pol II density (read counts/genebody length) in the genebody (TSS+500bp to CPS-500bp region).

Raw fastq sequencing files were uploaded to SRA under SRXXXXX accession ID. Processed data files were uploaded to GEO Database under the accession number GSEXXXXX.

### Comparison of EnChAMP-seq to CUT&RUN and ChIP-exo

Size fractionated, hg38 genome assembly mapped A549 cell line Pol II CUT&RUN data^75^ was downloaded from GEO Database (GSE155666). K562 cell line Pol II ChIP-exo data^76^ was downloaded from GEO Database (GSE108323). Genomic coordinates of the ChIP-exo data was converted from hg19 to hg38 using UCSC tools bigWigToBedGraph, liftOver, bedRemoveOverlap, and bedGraphToBigWig (v369)^135^. HCT116 PROcap corrected gene list was used for all heatmap and metagene profile plotting using DeepTools computeMatrix, plotHeatmap, and plotProfile (ver3.5.2)^130^.

### Processing of other external public data

HCT116 Pol II ChIP-seq data^136^ was downloaded from GEO Database (GSM5420207 and GSM5420209). Genomic coordinates of the ChIP-seq data was converted from hg19 to hg38 using UCSC tools bigWigToBedGraph, liftOver, bedRemoveOverlap, and bedGraphToBigWig (v369). Additional publicly available data used for this study were downloaded from ENCODE and GEO Databases include; HCT116 PROcap and PROseq data (GSE219427, GSE219376), HCT116 MNase data^137^ (GSE132705), HCT116 ATAC-seq data^138^ (GSM5904681 and GSM5904682). All of these data were processed similar to the CUT&RUN or the ChIP-exo data as described above.

### Visualization of Pol II structures

For the structural analysis of Pol II, we used representative structures from RCSB Protein Data Bank (PDB) for different transcriptional states, namely Pre-initiation^139^ (PDB ID# 8S55), Paused^21^ (PDB ID# 8UIS) and Elongation^6^ (PDB ID# 6GMH). Furthermore, full-length SUPT6H/SPT6 (UniProt ID# Q7KZ85) was modeled using AlphaFold3^140^ together with the Elongation complex. To depict the potential accessibility of GFP by the GFP-nanobody during the affinity purification experiments, the C-terminal region of the GFP was placed close to the N-terminal region of POLR2A/RPB1 using ChimeraX software^141^. The epitope regions on the GFP for the GFP-nanobody were annotated based on a published report^142^.

## Supplementary Tables

**Supplementary Table I.** List of all proteins detected in all EnChAMP-MS experiments used in this study. For each protein Gene name, pValue and log2Fold Change calculated from sample/GFP control, sample and GFP counts, and UNIPROT IDs are given. Note that each experiment, reported in separate sheets, involved 4 biological replicates of both the sample and the GFP control samples.

**Supplementary Table II.** Sequence of sgRNAs and homology arm oligos used for CRISPR/Cas9 gene editing of POLR2A/RPB1 and NELFA genes, as well as genotyping oligos used for verification of gene editing.

## Supplementary Data

**Supplementary Data I.** Complete sequence of HDR template constructs used for CRISPR/Cas9 engineering of GFP-control, GFP-RPB1, and NELFA-GFP endogenously tagged cell lines. SnapGene (.dna format) files are zipped together into a single file.

**Supplementary Data II.** Complete gene list, derived from NCBI RefSeq curated genes with PROcap corrected TSS coordinates that are expressed in HCT116 cells, used for all EnChAMP-seq analysis in bed format.

## Author Contributions

AO, HY, and JTL designed the project and secured the funding. MDD generated GFP control, GFP-RPB1, and NELFA-GFP cell lines. JJK, YT, and KO performed EnChAMP experiments, JJK, YT, YK, and SY performed mass spectrometry and data analysis for EnChAMP-MS, AO performed library preparation and data analysis for EnChAMP-seq. AO wrote the manuscript with the help from all the authors.

## Acknowledgments

Authors thank the members of the Lis and Yu labs, as well as the other co-PIs/co-Is Eftychia Apostolou, Steven Z. Josefowicz, Elizabeth Kellogg, Thomas Graham, William K. Lai, Hennig Lin, and the members of their labs, for discussion and critical insights during the project. We thank the Cornell University Epigenomics Core Facility for sequencing, Cornell Institute of Biotechnology Flow Cytometry Facility for help with cell sorting. We thank James Borovilas for help with the structural analysis. This work was supported by funding from NIH to AO, HY, and JTL (RM1 GM139738).

Authors declare no competing interest.

## Figure Legends

**Supplementary Figure 1. Quality control of EnChAMP.** A) Proper biochemical fractionation of cellular components during the steps of EnChAMP were verified with Western Blot analysis. Tubulin, Actin, and GAPDH are cytoplasmic marker proteins, and Histone 3 (H3) serves as the nuclear/chromatin-bound marker protein. B) Venn diagram showing the overlap between GFP-RPB1 interactor proteins identified in 3 independent GFP-RPB1 EnChAMP-MS experiments each carried out with 4 biological replicates of GFP-RPB1 and GFP control cell line samples. C) Scatterplot of log2(Fold Change) of GFP-RPB1 interactors identified in 3 replicate experiments. Pairwise comparison between Rep1-Rep2 (left), Rep1-Rep3 (middle), and Rep2-Rep3 (right panel) experiments. All common interactors between the compared experiments are plotted. Interactors identified in all three experiments are colored red, otherwise colored gray. Pearson correlation coefficient r is shown in each plot. D) Coverage of a 28Mb random region on Chromosome 1 in EnChAMP-seq input samples (prior to anti-GFP nanobody pull-down). GFP control Input in green, GFP-RPB1 Input in blue, GFP control +NVP2 Input in darkgreen, and GFP-RPB1 +NVP2 Input in darkblue. Genes within this region are shown below the tracks. E) Coverage of GAPDH gene (TSS-1Kb to CPS+1Kb) in GFP control (green), GFP-RPB1 (blue), GFP control +NVP2 (darkgreen), and GFP-RPB1 +NVP2 (darkblue) EnChAMP-seq samples. Gene annotations are shown below the tracks with blue rectangles.

**Supplementary Figure 2. Pol II interactors identified in other interactome studies.** A) Pol II interactors identified by POLR2A/RPB1 pull-downs in BioPlex, Huttlin *et.al.*, Cell, 2021. B) Pol II interactors identified by POLR2A/RPB1 pull-downs in OpenCell, Cho *et.al.*, Science, 2022. C) Pol II interactors identified by POLR2C/RPB3 pull-downs in Baluapuri *et.al.*, Mol. Cell, 2019. D) Comparison of identified Pol II interactor complexes’ completeness between EnChAMP, this study, and Baluapuri *et.al.*, Mol. Cell, 2019.

**Supplementary Figure 3. Distribution of EnChAMP-seq reads near TSS under drug treatment conditions.** Genome browser snapshot of reads mapped near the TSS of GAPDH gene under -NVP2 (left panel), +NVP2 (second from left panel), -Triptolide (second from right panel), and +Triptolide (right panel). The red dashed lines mark the PROcap detected major TSS of GAPDH gene, whereas the gray dashed lines mark the +/-150bp from TSS. For each of the drug treatments the matching no drug, -NVP2 and -Triptolide, samples are shown.

**Supplementary Figure 4. EnChAMP-seq determined Pol II DNA footprints.** Analysis of DNA footprint of RNA Pol II from GFP-RPB1 EnChAMP-seq with and without NVP2 treatment, replicate 1 (A), replicate 2 (B), with and without Triptolide treatment, replicate 1 (C), replicate 2 (D), and from NELFA-GFP EnChAMP-seq with and without NVP2 treatment (E). In each plot, read start coordinate relative to PROcap determined gene TSS is plotted on x-axis, while the read length is plotted on the y-axis. Cumulative readcounts from all HCT116 expressed genes were plotted with the aforementioned coordinates. Gray color scale is used to indicate cumulative readcounts at each position. Blue arrows mark the PIC footprints, while the red arrows mark the paused Pol II footprint.

**Supplementary Figure 5. Available structures of PIC, paused, and elongating Pol II complexes.** The N-terminal GFP tag on POLR2A/RPB1 is shown in cyan with red residues highlighting the epitope recognized by the anti-GFP nanobody. RPB1, the largest subunit of RNA Polymerase II is shown in gold, while the other Pol II subunits are colored in gray. Full-length SUPT6H/SPT6 modelled by AlphaFold3 in the elongation complex is shown in purple. Left panel: PIC (PDB ID# 8S55), Middle panel: Paused Complex (PDB ID# 8UIS), and Right panel: Elongation Complex (PDB ID# 6GMH with AlphaFold3 modeled SPT6).

